# Optical Pooled Screening for the Discovery of Regulators of the Alternative Lengthening of Telomeres Pathway

**DOI:** 10.1101/2025.02.15.638448

**Authors:** Isabel Quintanilla, Benura Azeroglu, Md Abdul Kader Sagar, Travis H. Stracker, Eros Lazzerini Denchi, Gianluca Pegoraro

## Abstract

Telomere elongation is essential for the proliferation of cancer cells. Telomere length control is achieved by either the activation of the telomerase enzyme or the recombination-based Alternative Lengthening of Telomeres (ALT) pathway. ALT is active in about 10-15% of human cancers, but its molecular underpinnings remain poorly understood, preventing the discovery of potential novel therapeutic targets. Pooled CRISPR-based functional genomic screens enable the unbiased discovery of molecular factors involved in cancer biology. Recently, Optical Pooled Screens (OPS) have significantly extended the capabilities of pooled functional genomics screens to enable sensitive imaging-based readouts at the single cell level and large scale. To gain a better understanding of the ALT pathway, we developed a novel OPS assay that employs telomeric native DNA FISH (nFISH) as an optical quantitative readout to measure ALT activity. The assay uses standard OPS protocols for library preparation and sequencing. As a critical element, an optimized nFISH protocol is performed before in situ sequencing to maximize the assay performance. We show that the modified nFISH protocol faithfully detects changes in ALT activity upon CRISPR knock-out (KO) of the *FANCM* and *BLM* genes which were previously implicated in ALT. Overall, the OPS-nFISH assay is a reliable method that can provide deep insights into the ALT pathway in a high-throughput format.

## 1. Introduction

Replicative immortalization via constitutive elongation of telomeres in somatic cells is one of the key hallmarks of cancer [1]. In about 90% of cancer cell types, the telomerase enzyme (TERT) is reactivated [2]. In the remaining 10-15%, including several osteosarcomas, replicative immortalization, telomere elongation is instead achieved by a telomerase-independent mechanism, referred to as Alternative Lengthening of Telomeres (ALT). ALT employs a type of homologous recombination, break-induced replication, to elongate telomeres [3,4]. The ALT pathway represents a promising therapeutic target in this subclass of tumors because it can be identified cytologically and genetically. Targeting ALT could be useful due to both direct effects on telomere lengthening, that is essential for replicative immortalization in cancer cells, and indirect effects, as inhibition of other cellular pathways might be synthetically lethal with ALT. Thus, the identification of cellular factors that are either necessary for ALT, or that are lethal in ALT cells when inhibited, represents an important anti-cancer target identification strategy [5,6].

The identification of ALT targets depends on cellular assays that measure ALT activity in cancer cells. Recent work from our laboratories showed that ALT cells display a high level of telomeric single-stranded DNA (ssDNA) when compared to non-ALT cells, which can be detected by native DNA FISH (nFISH) [7]. Furthermore, we demonstrated that nFISH can be miniaturized and automated in 384-well plates to measure changes in ALT activity levels upon CRISPR-Cas9 knock-out (KO) of known cellular factors [8]. Using this assay, we screened a library containing more than 1,000 CRISPR-Cas9 KO reagents in an arrayed format to identify genes whose KO results in either inhibition or increase of ALT activity [9]. Despite these recent advances, there are still significant limitations regarding the throughput and the type of functional perturbations (e.g. CRISPR-KO, CRISPR interference, or CRISPR activation) that can be evaluated in arrayed format.

Optical pooled screens (OPS) have recently emerged as a powerful alternative to overcome the limits of arrayed functional genomics assays with imaging readouts [10,11]. This approach combines pooled lentiviral libraries with *in situ* sequencing (ISS) to relate genetic perturbations to visual phenotypes at the single cell level. Importantly, while OPS assays have been used for functional genomics screens paired with fluorescent dyes, immunofluorescence, and RNA FISH as readouts [10,12–15], DNA FISH assays, such as nFISH, have not been yet used in conjunction with OPS, possibly due to technical challenges. To fill this gap, and to extend the throughput and range of CRISPR-based functional genomics perturbations that can be implemented to identify ALT genes, we describe here the optimization of an OPS assay with telomeric nFISH as a readout.

## 2. Results

### 2.1. Implementation and optimization of Optical Pooled Screens (OPS)

To optimize the previously described OPS protocols [10,11] in our experimental setup, we used three different non-transformed and cancer cell lines with constitutive expression of *S. pyogenes* Cas9 (spCas9): hTERT-RPE-Cas9, HCT116-Cas9, and U2OS-Cas9. sgRNA expression was achieved by transducing cell lines at a low multiplicity of infection (MOI < 0.1) with an equimolar mix of five sgRNAs: three targeting the *LMNA* gene (sgLMNA), encoding for lamin A (a major component of the nuclear lamina), and two serving as non-targeting negative control (sgCTL). Following transduction, cells were selected with puromycin to isolate cells expressing both Cas9 and sgRNA (Fig. 1A). The heterogenous population of cells was fixed and processed according to the standard OPS protocol [10,11] for barcode library preparation and *in situ*-sequencing (ISS) using a 4-color sequencing by synthesis (SBS) chemistry (Fig. 1A). Cells were subjected to 12 cycles of SBS, each involving fluorophore incorporation, imaging, and fluorophore cleavage, using a 10X objective on an automated widefield microscope (Fig. 1A and 1B; see Materials and Methods for details). An adapted image bioinformatics pipeline [11] was used to segment nuclei based on DAPI staining. All images were aligned across all cycles using the DAPI channel and the fluorescence intensity in 4 channels was used to assign ISS spots to single cells (See Materials and Methods and for details). Using this analysis pipeline, 4.75 ± 1.18, 4.17 ± 0.9 and 4.34 ± 1.6 ISS spots per cell were detected for hTERT-RPE-Cas9, HCT116-Cas9 and U2OS-Cas9 cells, respectively, expressing a mix of 5 sgRNAs (mean ± SD, Fig. 1B and 1C). Of note, ISS spots were only quantified when they were consistently detected across all 12 cycles. Base calling analysis revealed that approximately 80% of sequence reads mapped to one of the expected sgRNA sequences for all cell lines. As expected, we observed that genotyping accuracy was inversely related to the number of sgRNA bases sequenced and directly related to spot quality (Fig. 1D and 1E). As SBS is based on iterative rounds of fluorophore incorporation and cleavage, a steady decrease in spot quality and an increase in fluorescence background is to be expected but manageable with additional washes.

**Figure 1.**
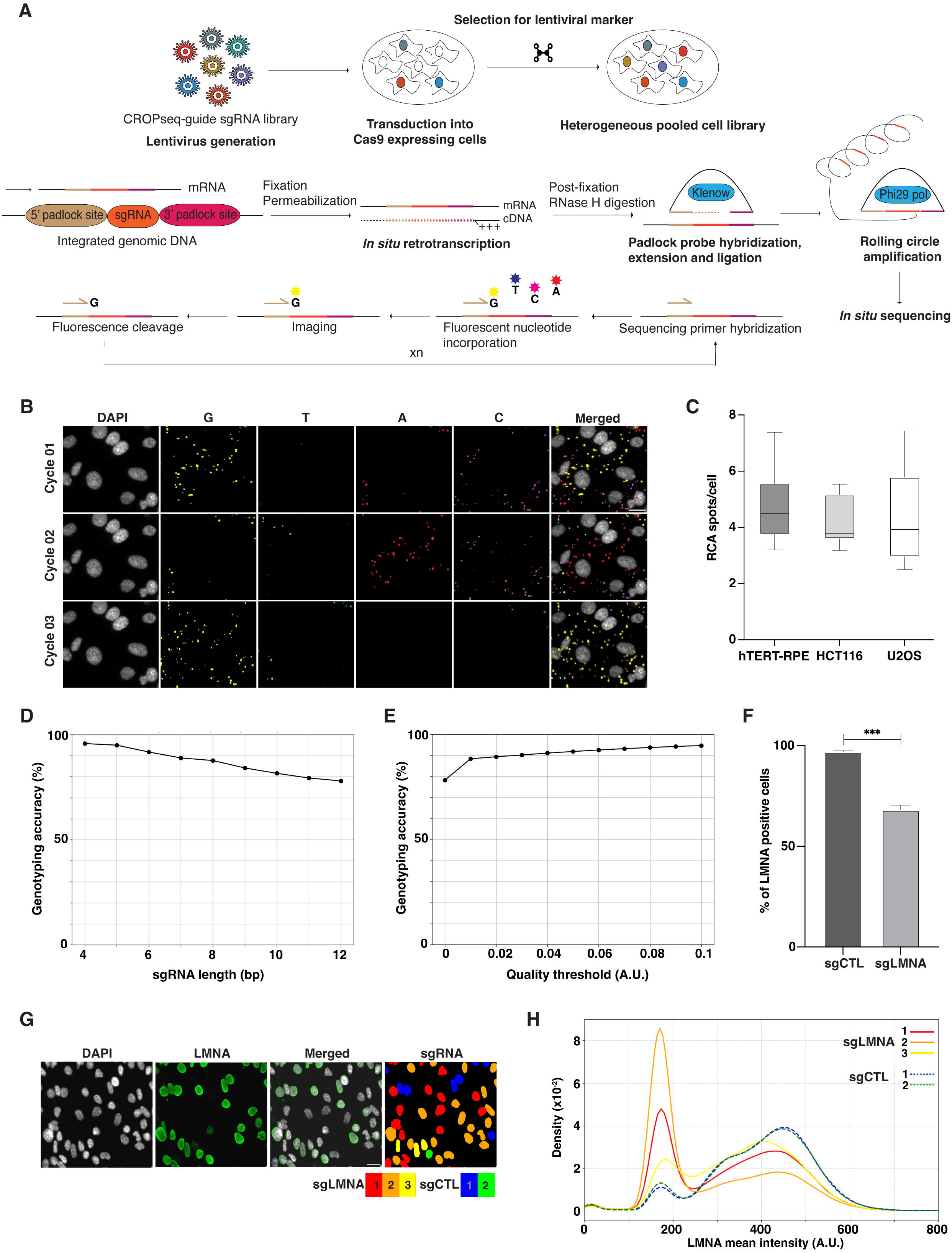
Overview of the OPS workflow and its implementation. **A)** OPS experimental workflow. A lentiviral library containing sgRNAs of interest was transduced into target spCas9-expressing cells at a low multiplicity of infection (MOI), followed by selection for a resistance marker to generate a heterogenous pooled cell library. Cells with an integrated lentiviral vector were then fixed and permeabilized. Subsequently, the mRNA containing the sgRNA sequence was retrotranscribed *in situ* using an LNA-modified oligo. Next, a padlock probe was hybridized around the sgRNA sequence of the cDNA followed by probe extension and ligation. The circularized probe containing the sgRNA sequence was used as a template to perform rolling circle amplification (RCA) and the resulting amplification product is sequenced using sequencing-by-synthesis (SBS) chemistry, a form of *in situ* sequencing (ISS). **B)** Representative images of 3 ISS cycles using 4-color SBS chemistry. Scale bar: 20 microns. **C)** Boxplots showing the number of RCA spots per cell detected in three different cell lines expressing 5 sgRNAs at a time and using conventional OPS protocol. The middle line represents the 50^th^ percentile. The edges of the box represent the 25^th^ and 75^th^ percentile, respectively. The whiskers extend to 1.5 * Inter Quantile Range (IQR). **D)** Plot showing the inverse relationship between the length of the sgRNA read and genotyping accuracy when using ISS. **E)** Graph illustrating the direct relationship between spot quality threshold and genotyping accuracy when using ISS. **F)** Bar plot showing the percentage of lamin A-positive cells when hTERT-RPE-Cas9 cells are transduced with just a control non-targeting sgRNA (sgCTL) or an equimolar mix of 3 *LMNA*-targeting and 2 scrambled/non-targeting sgRNAs (sgLMNA). Mean± SD. **G)** Example images of hTERT-RPE-Cas9-sgLMNA cells stained with an anti-lamin A primary antibody and a Alexa488-conjugated secondary antibody together with a segmentation and sgRNA classification mask. Scale bar, 20 microns. **H)** Density plot showing the distribution of lamin A mean intensity values across hTERT-RPE-Cas9-sgLMNA cells. The proportion of cells with lower lamin A intensity values is larger in cells expressing *LMNA*-targeting sgRNAs in comparison to cells expressing non-targeting sgRNAs.

Next, we sought to measure the degree of association between individual sgRNA, and a phenotype measured by imaging. To this end, we used the hTERT-RPE-Cas9 cell line expressing five sgRNAs (including *LMNA*-targeting and control non-targeting). Cells were first processed for OPS library preparation for up to and including the post-fixation step (Fig. 1A), and then for immunofluorescence (IF) with an antibody against lamin A. Cells were then imaged with a 20X objective, and lamin A fluorescence levels at the single cell level were measured using automated image analysis (see Materials and Methods for details). Using this approach, we observed a reduction in lamin A staining of approximately 30% in the pool of cell expressing a sgRNAs against lamin A (sgLMNA + sgCTL) compared to the control sample (sgCTL) (Fig. 1F). After IF, we finished the OPS library preparation and processed the sgLMNA + sgCTL pool of cells for 12 cycles of ISS and imaging at 10X, we then ran the ISS bioinformatics analysis pipeline on them, and we finally computationally matched cells at different magnifications in the ISS and IF images, respectively (See Materials and Methods for details), to match sgRNA sequences with lamin A IF intensity values at the single cell level (Fig. 1G and 1H). As a result, we observed that lamin A IF intensity values were substantially lower in cells expressing 2 out of the 3 sgRNAs targeting the *LMNA* gene when compared to cells expressing non-targeting control sgRNAs, whereas one LMNA sgRNA resulted in only a modest knockdown. These results establish a robust workflow for CRISPR-KO in a pooled population of cells.

### 2.2. Optimization of telomeric ssDNA native FISH (nFISH) with In Situ Sequencing

When integrating fixed-cell phenotyping with ISS, it is essential to properly combine protocols to allow for phenotype detection while preserving the integrity of the sgRNA sequence expressed as an mRNA. This is because some phenotype-labelling protocols can lead to RNA degradation, and at the same time some steps in the ISS protocol might compromise phenotype detection. To maximize OPS performance, phenotyping steps should be performed either after the retrotranscription (RT) and post-fixation steps or prior to the SBS cycles (Fig. 2A and [11]). Before attempting to combine telomeric nFISH and ISS, we assessed both methods individually in ALT-positive U2OS cells [7] stably transduced with a lentiviral vector expressing a control non-targeting sgRNA.

**Figure 2.**
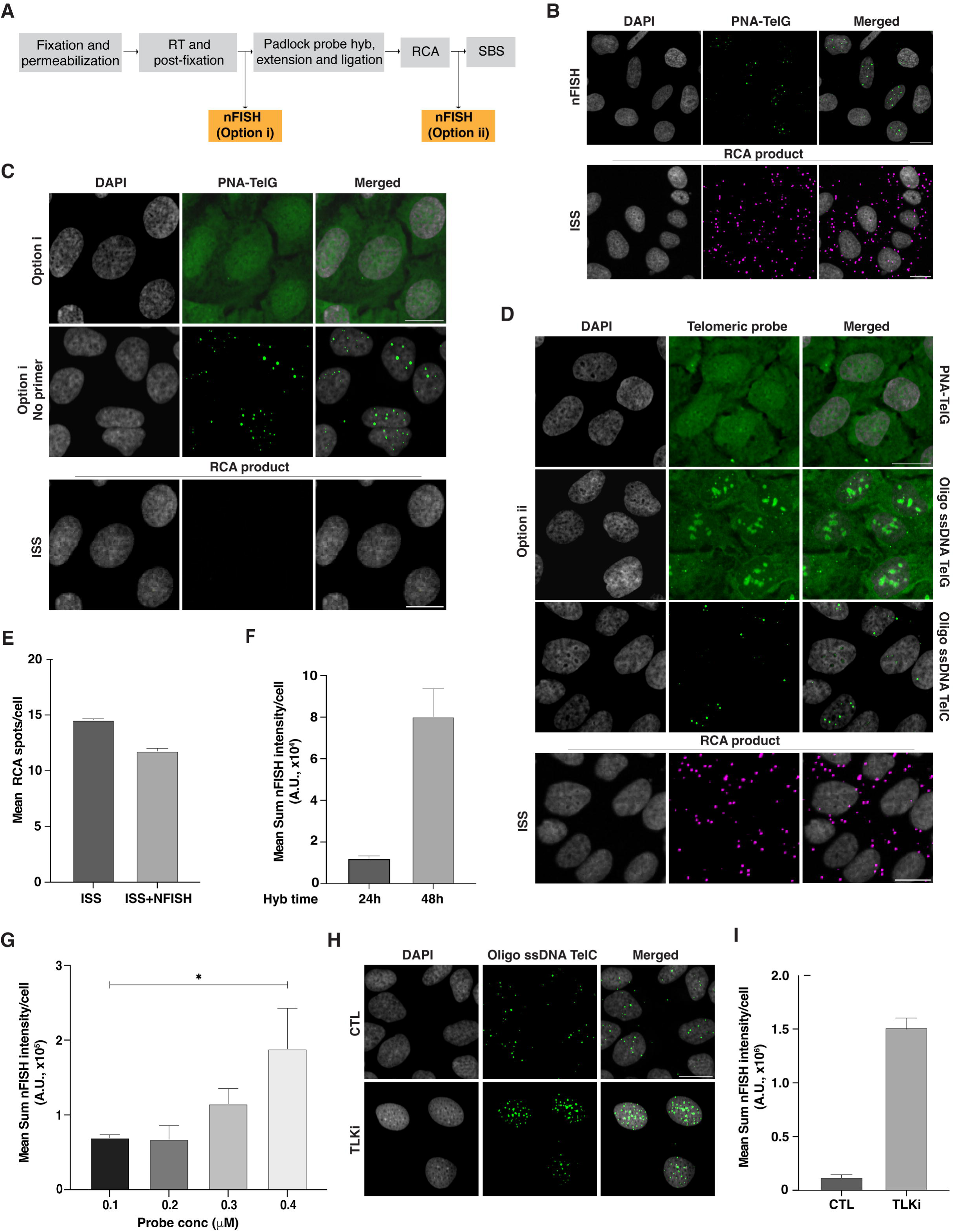
Optimization of the OPS-nFISH protocol and its integration with ISS. **A)** OPS experimental workflow illustrating the steps in the ISS library preparation where the nFISH protocol can be integrated. **B)** Example nFISH and ISS images when both protocols are performed individually in U2OS-Cas9-sgScramble cells. ISS/RCA spot images are obtained after hybridizing a fluorescent probe against a constant region of the sgRNA-padlock probe construct previously amplified by RCA. Scale bar: 20 microns. **C)** Representative nFISH and ISS images when the phenotyping protocol is performed after retrotranscription (RT) and post-fixation (Option i) in U2OS-Cas9-sgScramble cells. (Top panel) nFISH images when a PNA-TelG probe is used. (Middle panel) nFISH images when RT primer is removed from reaction. (Bottom panel) ISS images showing the lack of RCA spots when ISS and nFISH are combined using Option i. Scale bar: 20 microns. **D)** Representative nFISH and ISS images when the phenotyping protocol is performed after RCA reaction (Option ii) and using different telomeric probes in U2OS-Cas9 cells transduced with sgCTL. (Top panel) nFISH images when the PNA-TelG probe is used. (Middle panels) nFISH images when oligo ssDNA TelG or oligo ssDNA TelC are used. (Bottom panel) Example RCA spot images regardless of telomeric probe used for nFISH. Scale bar: 20 microns. **E)** Bar plot of the difference in mean RCA spots per cell between ISS protocol performed individually or in combination with nFISH using Option ii. Mean± SD. **F)** Plot showing the difference in telomeric integrated fluorescence intensity between 24 h and 48 h of oligo ssDNA TelC probe hybridization. Mean± SD. **G)** Bar plot illustrating how increasing the oligo ssDNA TelC probe concentration results in a steady increase in telomeric integrated fluorescence intensity. Mean± SD. **H)** Example nFISH images showing the increase in telomeric signal after U2OS-Cas9-sgScamble cells are treated with 10 mM of a TLK inhibitor (TLKi) for 6 h. Scale bar: 20 microns. **I)** Quantification of telomeric integrated fluorescence intensity for non-treated and TLKi-treated U2OS-Cas9-sgScramble cells. Mean± SD.

As expected and previously reported, performing telomeric nFISH alone in U2OS cells in the absence of ISS resulted in discrete spots inside the nucleus corresponding to telomeric ssDNA (Fig. 2B; [8]). Similarly, we could successfully detect the formation of the ISS substrate when performing the ISS protocol under FISH conditions, as shown by the appearance of several spots in cells FISH-stained with a fluorescently labelled oligo targeting a constant region of the sgRNA-padlock probe construct after ISS library preparation (Fig. 2B). Having validated nFISH and ISS protocols independently in U2OS cells, we next attempted to integrate telomeric nFISH by performing it after the RT and post-fixation steps of the ISS protocol (Fig. 2A, option i). Unexpectedly, this approach resulted in high nuclear fluorescence background and in the absence of specific telomeric spot-like signals in the nFISH channel (Fig. 2C). Systematic elimination of individual RT components indicated the chemically modified RT primer as the cause of the high fluorescence background. Removing just this component from the protocol completely suppressed the fluorescence background and restored the telomeric spot-like signals in the nFISH channel (Fig. 2C, middle panel). However, several attempts at optimization of this step, including (i) hybridization of the chemically modified primer before the RT reaction followed by intense washes to remove non-specific binding of the primer, (ii) decreasing the RT primer concentration, or (iii) increasing the number and stringency of the washes after the RT reaction did not improve the fluorescent background in the nFISH channel. Furthermore, and more importantly, we also observed that performing the telomeric nFISH after RT and post-fixation compromised the efficiency of rolling circle amplification (RCA) product formation during the protocol, which is the main substrate for ISS (Fig. 2C). We thus concluded that performing nFISH under these conditions was not compatible with ISS and OPS.

To overcome this issue, we performed telomeric nFISH after the RCA step, and right before the SBS cycles (Fig 2A, option ii). However, in these experimental conditions, the telomeric nFISH channel continued to show high fluorescence background (Fig. 2D), suggesting the need for additional changes to the nFISH protocol. Accordingly, we applied several modifications to the standard nFISH protocol [8], including switching from the fluorescently labelled TelG PNA oligo probe to fluorescently labelled TelG or TelC ssDNA oligo probes (here by referred to as unmodified ssDNA oligo probes). In addition, to accommodate the use of ssDNA oligo probes, the nFISH hybridization buffer was modified to reduce the formamide content from 70% to 50%, and the blocking buffer content from 0.5% to 0.2%. These adjustments account for the lower binding affinity of the unmodified ssDNA oligo probes when compared with the PNA probes. For the same reason, the nFISH probe hybridization time was increased from 1 h to 24 h, and the post-nFISH hybridization washing buffer composition was changed to remove formamide and replace it with 2X SSC and Tween-20 instead. Using this modified protocol and the TelG ssDNA oligo probe, the nFISH staining resulted in the expected spot-like telomeric signal, but also in high non-specific signal likely inside the nucleolus and in high nuclear fluorescence background. In similar experimental conditions, instead, using the TelC ssDNA oligo probe resulted in a slightly reduced but spot-like telomeric signal pattern, with minimal nucleolar or nucleoplasmic background, resembling the staining pattern observed when performing nFISH staining in isolation (Fig. 2D and Fig. 2A). Additionally, using the option ii protocol (telomeric nFISH between the RCA step and the first SBS cycle) resulted in only minimally reduced the number of RCA products available for ISS, regardless of the telomeric ssDNA oligo probe used for nFISH (14.55 ± 0.1 vs 11.77 ± 0.24 RCA spots/cell, mean± SD) (Fig. 2D and Fig. 2E).

To improve the detection efficiency of telomeric nFISH for ISS using the TelC ssDNA oligo probe, we further optimized the hybridization time and probe concentration. We observed the integrated fluorescence intensity of the spot-like telomeric nFISH signal was increased 6.7-fold when the probe was hybridized for 48 h compared to 24 h (Fig. 2F). Also, the telomeric nFISH fluorescence signal was further enhanced by a 4-fold increase in the probe concentration during the hybridization reaction (Fig. 2F and Fig. 2G). Finally, we assessed whether the modified nFISH protocol with the TelC ssDNA oligo probe retained specificity for telomeric ssDNA, and whether it could be used to quantify changes in ALT activity as measured by a change in nFISH signal. To this end, we treated U2OS cells with a chemical inhibitor, E804-20 [16] (referred to henceforth as TLKi), of the Tousled Like Kinase 1 and 2 proteins (TLK1 and TLK2), which are involved in regulation of chromatin assembly [17], and whose chemical inhibition had been previously shown to lead to an increase in the telomeric nFISH signal [9]. As expected, we observed a 12-fold increase in telomeric nFISH signal after treating the U2OS cells with 10 μM of TLKi for 6h and processing them using the TelC ssDNA oligo protocol (Fig. 2H).

Taken together, these results indicate that the nFISH step must be performed right before the first SBS cycle to successfully integrate telomeric nFISH with the ISS protocol. Furthermore, they show that an optimized nFISH protocol using TelC ssDNA oligo probe is required for compatibility with ISS. These findings validate the optimized nFISH protocol for the specific detection of telomeric ssDNA, and its robustness in detecting changes ALT-associated telomeric ssDNA levels upon a known experimental perturbation.

### 2.3. CRISPR-KO of known ALT-regulating genes in a pooled lentiviral format leads to an increased nFISH signal

To test for differential telomeric nFISH detection following the knockout of a known ALT regulator, we transduced stable U2OS-Cas9 cells with a pool of 4 lentiviral vectors, each expressing a different sgRNA targeting the *FANCM* gene. *FANCM* encodes a member of the Fanconi Anemia complementation group, and is an essential gene involved in DNA repair. Notably, *FANCM* KO has been previously shown to increase the telomeric nFISH signal [9]. The transduction of U2OS-Cas9 cells with the pool of 4 sgRNA against *FANCM* resulted in cell death and a significant decrease in telomeric nFISH signal, rather than the expected signal increase, when compared to cells transduced with a lentiviral vector expressing a control non-targeting sgRNA (Fig. 3A). This is most likely due to the requirement of a minimum of 7-10 days to generate OPS cell libraries, which might lead to the loss of cells, where essential genes, including *FANCM* are targeted. Given that many potential ALT regulators may be essential [9], we reasoned that the stable U2OS-Cas9 cell line might not be a suitable genetic background to study ALT regulators using OPS.

**Figure 3.**
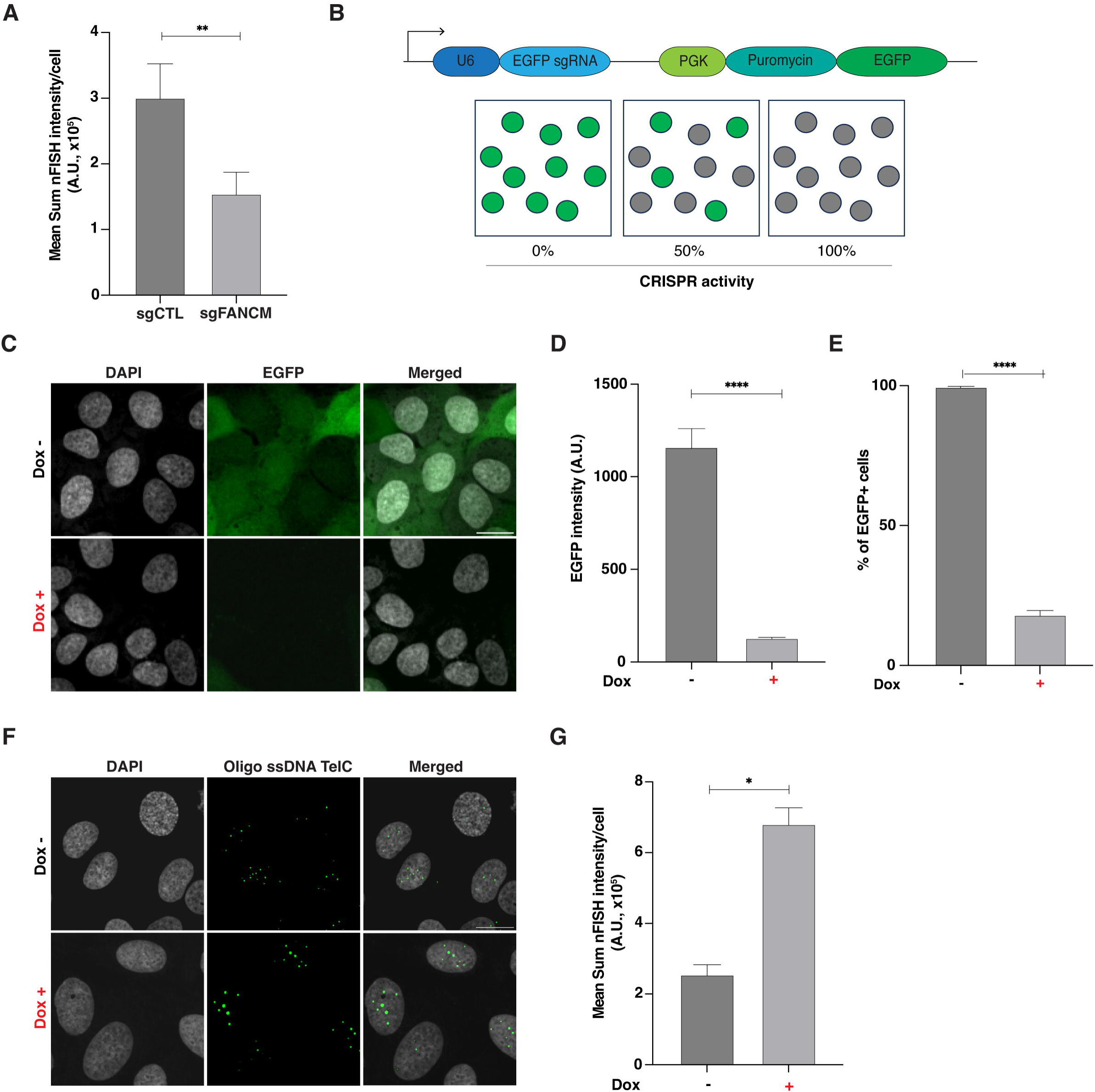
Validation of CRISPR efficiency and telomeric nFISH signal detection in U2OS-iCas9 cells. **A)** Quantification of the telomeric integrated fluorescence intensity for U2OS-Cas9 cells transduced with either lentiviral sgCTL alone or with an equimolar mix of 4 lentiviral *FANCM*-targeting and 2 different scrambled/non-targeting sgRNAs (sgFANCM). Mean± SD**. B)** Illustration of the lentiviral vector used to evaluate CRISPR efficiency, pXPR-011[29]. This vector expresses the gene *EGFP* and an EGFP-targeting sgRNA. **C)** Representative images of uninduced (Dox-) and induced (Dox+) U2OS-iCas9 cells transduced with pXPR-011. Scale bar: 20 microns. **D)** Bar plot showing the decrease in total EGFP intensity in induced (Dox+) U2OS-iCas9-pXPR-011 cells compared to uninduced (Dox-) cells. **E)** Graph showing a decrease in the percentage of EGFP-positive U2OS-iCas9-pXPR-011 cells when Cas9 expression is induced with doxycycline compared to uninduced cells. Mean± SD. **F)** Example nFISH images of uninduced (Dox-) and induced (Dox+) U2OS-iCas9 cells transduced with sgFANCM. Scale bar: 20 microns. **G)** Plot illustrating the increase in telomeric integrated fluorescence intensity for induced (Dox+) U2OS-iCas9-sgFANCM cells compared to uninduced (Dox-) cells. Mean± SD.

To identify essential genes involved in the ALT pathway using OPS, we generated a U2OS cell line with a doxycycline-inducible Cas9 system (U2OS-iCas9) and validated its activity using a lentiviral vector co-expressing the *EGFP* gene and a corresponding *EGFP*-targeting sgRNA (Fig. 3B). The results of this experiment showed that doxycycline (DOX) induction of Cas9 in U2OS-iCas9 cells resulted in a significant reduction of EGFP intensity and the percentage of EGFP-positive cells, corresponding to approximately 80% KO efficiency (1,200 vs 130 A.U. mean nFISH intensity/cell, and 99% vs 18% EGFP-positive cells, respectively, Figs. 3C-3E). Next, we generated a U2OS-iCas9 cell line expressing a pool of 4 *FANCM* sgRNAs and observed a significant increase (2.6-fold) in telomeric nFISH signal after a 3-day Cas9 induction when compared to the same uninduced pool of cells (Fig. 3F and 3G). Altogether, these results indicated that an inducible Cas9 system is necessary for studying essential genes using OPS. Furthermore, *FANCM* knockout in the U2OS-iCas9 cell line can be used as a positive biological control for functional genomics screens that measure changes in ALT activity.

### 2.4. Optical pooled screens enable accurate mapping of genotype-to-phenotype association for ALT regulators

Having optimized the OPS-nFISH assay and validated *FANCM* as a positive control, we next aimed to assess the ability of assay to detect changes in ALT activity. To do so, we used a set sgRNAs targeting either of the three genes involved in ALT activity, *FANCM*, *DDX39A* and *BLM*, as well as non-targeting control and a safe-targeting sgRNA, for a total of 14 sgRNAs (see Supplemental Material). *DDX39A* gene encodes a RNA helicase involved in transcriptional gene regulation and telomere maintenance in telomerase positive cells [18], whose KO has been described to increase ALT activity [9], while *BLM* gene encodes is a DNA helicase required for maintaining ALT activity by [19] and whose depletion is known to decrease telomeric nFISH signal [9,19]. We also used a “Safe” sgRNA, which is commonly used as controls in CRISPR-Cas9 screens and targets genomic regions not known to contain any genes and. To generate a pooled cell library, we transduced U2OS-iCas9 cell line at an MOI < 0.1 and processed it using the OPS-nFISH workflow (Fig. 4A). A total of 600,000 cells were imaged, and ISS spots were detected in 51% of those cells, with an average of 2.5 ISS spots per cell (Fig. 4B). After performing base calling analysis, 64% of sequence reads were successfully mapped to one of the expected sgRNA sequences and matched with corresponding nFISH signal to achieve genotype-to-phenotype association. Moreover, we observed that all the sgRNAs were present in the analyzed cell library, confirming proper cell representation (Fig. 4C). All sgRNAs targeting *FANCM* resulted in varying degrees of increased telomeric nFISH signal, with sgFANCM-1 producing the strongest effect and sgFANCM-3 the weakest, when compared to the sgCTL (3,316 vs. 1,144 A.U. mean nFISH intensity/cell, p-value < 0.0001 for sgFANCM-1 vs sgCTL and 1,362 vs. 1,144 A.U. mean nFISH intensity/cell, p-value < 0.0001 for sgFANCM-3 vs. sgCTL). Conversely, we did not observe any significant change in ALT activity in cells where *DDX39A* was targeted. The large number of cells imaged allowed us to also detect a discrete but significant decrease in telomeric nFISH signal for all the BLM-targeting sgRNAs. In this case, sgBLM-1 resulted in the strongest signal decrease and sgBLM-2 in the weakest, when compared to a control sgRNA (911 vs. 1,144 A.U. mean nFISH intensity/cell, p-value = 1.3 e-12 for sgBLM-1 vs. sgCTL and 967 vs 1,144 A.U. mean nFISH intensity/cell, p-value = 8.2 e-07 for sgBLM-2 vs. sgCTL) (Fig. 4D and E). Overall, these results demonstrate that the nFISH assay in an OPS format enables faithful genotype-to-phenotype associations and effectively identify genetic perturbations that have a range of effects on ALT biology, as even subtle changes in pathway activity are identified when imaging large numbers of cells.

**Figure 4.**
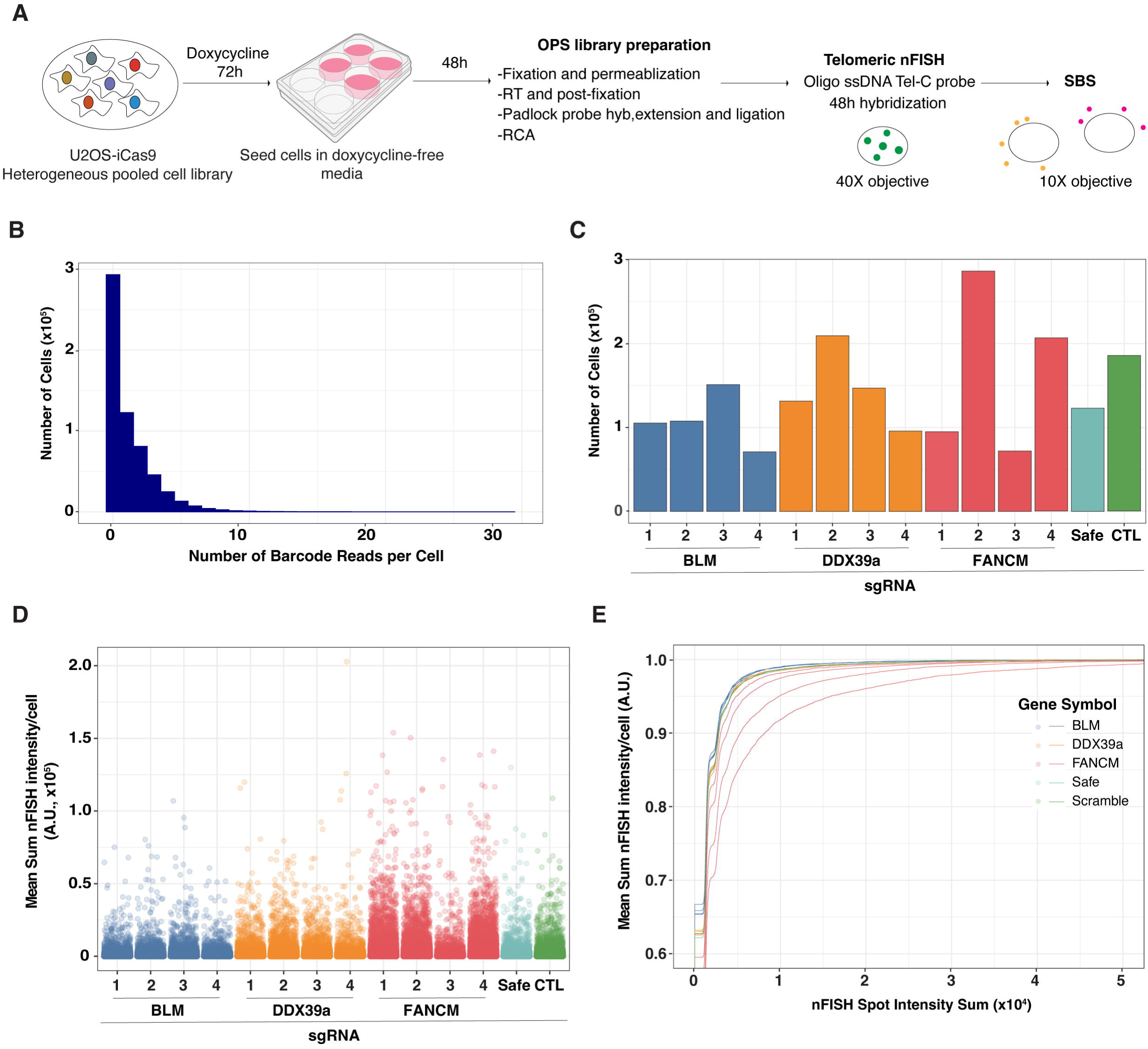
OPS-nFISH experimental workflow and analysis of telomeric signal detection. **A)** OPS-nFISH experimental workflow. spCas9 expression was induced in the pooled cell library by treating the cells with 0.05 μg/mL doxycycline for 72h followed by seeding the cells in a 6-well plate using doxycycline-free media. After 48h, cells were fixed and processed for OPS library preparation. Then, telomeric nFISH was performed using the modified oligo ssDNA protocol, followed by SBS reactions to read the sgRNAs expressed in each cell. **B)** Plot showing the number of sgRNA barcode reads per cell detected during ISS in the pooled cell library. **C)** Bar plot showing the number of cells expressing each sgRNA in the library. **D)** Single-cell quantification of telomeric integrated fluorescence intensity associated to each sgRNA. **E)** Empirical Cumulative Distribution Function (ECDF) plot showing the cumulative distribution of telomeric integrated fluorescence intensity based on the different sgRNAs present in the library.

## 3. Discussion

The identification of the molecular mechanisms underlying tumor initiation and progression is crucial for advancing our understanding of cancer and developing novel oncology therapeutic strategies. The ALT pathway of telomere elongation is an attractive target for cancer therapy but remains poorly characterized. A major challenge in studying ALT has been the lack of scalable detection methods suitable for high throughput screening (HTS) to identify critical players and regulators of this pathway. One of the most commonly used methods to detect ALT, the C-circle assay, which is the gold standard to measure ALT activity [20], requires isolation of genomic DNA and uses RCA to detect the extrachromosomal telomeric repeats (ECTR) that are associated with this pathway. Unfortunately, this method provides population-based information and has limited throughput [20] and as such, is not suitable for use in large-scale genomic screening approaches. An alternative method for detection of ALT is the imaging-based telomeric nFISH which was used to measure single-stranded telomeric DNA associated with ECTR [8]. nFISH is a robust, quantitative and imaging-based assay to measure ALT activity at a single cell level that can be applied to arrayed functional genomic screens [8,9,19,21,22]. While arrayed HTS enables a straightforward genotype-to-phenotype association, it is time-consuming, requires specific liquid handling equipment, and is limited the scale and types of genomic perturbations that can be studied. Conversely, pooled screens with optical readouts offer a highly flexible, scalable, cost- and time-effective alternative that significantly extents discovery potential of HTS [10,11,23]. Therefore, integrating telomeric nFISH with OPS represents a powerful approach for advancing breakthrough discoveries in ALT biology.

To address this technological need, we developed a novel OPS assay pipeline that uses telomeric nFISH as its optical readout for studying ALT. We found that, to successfully integrate telomeric nFISH with ISS, the phenotypic assay needs to be performed right before the first SBS cycle and after the RCA reaction, the final step of the ISS library preparation. Moreover, since the conventional telomeric nFISH assay was not compatible with OPS, we modified and validated the nFISH protocol to allow for the use of oligo ssDNA telomeric probes, whose performance remains unaffected by the ISS reactions.

Since its implementation OPS has contributed to advances in various fields, including viral infection, immune response, and cell morphology [10,12–14,24,25]. However, the imaging assays used in these studies have not explored the use of DNA FISH. This is likely because conventional OPS relies on detecting the sgRNA sequence as a part of an mRNA transcribed by RNA pol II, which can be compromised by certain phenotyping assays [11], such as DNA FISH. Integrating DNA FISH with OPS represents a technical challenge due several factors: the high temperatures required to denature genomic DNA, the use of RNAse in certain DNA FISH protocols, and the need for high-resolution microscopy to accurately capture DNA FISH fluorescent signals. In contrast, the telomeric nFISH assay used in this study detects ssDNA, eliminating the need for DNA denaturation, and the resulting spot-like fluorescence signal can be reliably detected with a 40X objective, making the assay easily scalable. Unfortunately, though, our first assay development experiments revealed that when telomeric nFISH was performed after the RT and post-fixation steps, a common approach when pairing phenotypic assays with ISS [11], it impaired the proper formation of RCA products, which are essential for cell library genotyping. Furthermore, we observed that the RT primer compromised the performance of the telomeric nFISH when the gold-standard PNA probe was used. Together, these findings highlights the necessity of optimizing each combination of phenotyping and ISS assay to achieve the best performance of both methods. [11] We solved this problem by performing the nFISH protocol after the RCA products are formed, therefore preventing any interference between the phenotyping protocol and the sequencing library preparation, while the change in telomeric probe chemistry resolved the fluorescence background issues caused by the RT primer.

We demonstrate here that OPS-nFISH can accurately detect genetic perturbations that increase the ALT activity and corresponding telomeric nFISH signal (Fig 4). By enabling large-scale studies of ALT biology and accommodating a wide range of genetic perturbation tools, such as CRISPRa and CRISPRi, OPS expands the ability to identify key regulators beyond those detectable with the conventional CRISPR-KO approach. Moreover, it creates opportunities to perform follow-up base editor screens, allowing precise characterized of functional variants of previously identified genetic targets [26]. Overall, this method provides a comprehensive framework to gain insightful understanding into the ALT pathway.

The main limitation of the OPS assay in its current implementation is that the baseline fluorescence nFISH telomeric signal is low, even in ALT-positive U2OS cells. This reduces the chance of reliably identifying CRISPR-KO treatments that result in a reduction of the nFISH signal and would potentially require imaging of a larger number of cells, significantly increasing the time needed to perform a screen. When the *in situ* detection steps are known to hinder the efficiency of the phenotyping assay, performing the phenotyping assay before the ISS library preparation is recommended [11]. However, in our case, the RNAse A treatment in the nFISH protocol prevented this option. To address this technical shortcoming, a T7 promoter has recently been incorporated into the OPS lentiviral vector, enabling *in situ and in vitro* transcription of sgRNAs after the phenotyping assay and before the ISS steps [15,27]. Such methodology improvement enables to phenotype the cell library before starting the ISS library preparation steps, which enhances phenotyping performance and allows the use of highly multiplexed phenotypic readouts, and to better control the transcription levels of the sgRNA leading to greater yield of sequencing products. Therefore, integrating telomeric nFISH with the novel T7-OPS approach could further improve the results described in this study.

In summary, the OPS-nFISH method described here has the potential to expand the range and the throughput of functional genomics screens towards deeper understanding of the molecular basis of ALT, and to advance the discovery of potential ALT-specific therapeutic targets.

## 4. Materials and Methods

### 4.1. Tissue culture

hTERT-RPE-Cas9 [28] and U2OS-Cas9 cells were cultured in High-glucose DMEM (ThermoFisher Scientific #11965-092) supplemented with 2 mM L-Glutamine (ThermoFisher Scientific #25030-081), 100 U/mL penicillin-streptomycin (ThermoFisher Scientific #15140-122), 10% heat-inactivated fetal bovine serum (ThermoFisher Scientific #10082-147), and 10 μg/mL Blasticidin (Gibco #A1113903). U2OS-iCas9 cells were cultured in the same media with serum substituted for 10% tetracycline-screened fetal bovine serum (Takara #631101) and antibiotic selection substituted for 800 μg/mL Geneticin (ThermoFisher Scientific #10131-027). HCT116-Cas9 [29] cells were cultured in RPMI (ThermoFisher Scientific #11875093) supplemented with 2 mM L-Glutamine, 100 U/mL penicillin-streptomycin, 10% heat-inactivated fetal bovine serum, and 4 μg/mL Blasticidin. HEK293FT cells were cultured in High-glucose DMEM supplemented with 2 mM L-Glutamine, 100 U/mL penicillin-streptomycin, 10% heat-inactivated fetal bovine serum, and 1mM sodium pyruvate (ThermoFisher Scientific #11360-070).

### 4.2. Stable and Inducible Cas9 cell line generation

The U2OS-Cas9 cell line was generated by transducing U2OS (ATCC HTB-96) cells with the lentiviral vector pLX311-Cas9 (Addgene #118018). After transduction, cells were selected with 10 μg/mL Blasticidin and single clones were generated using limiting dilution. The most optimal clone was selected based on Cas9 expression levels quantified by immunofluorescence.

The U2OS-iCas9 cell line was generated by transfecting U2OS cells with a *piggyBac* transposase plasmid [30] and the HP138-neo *piggyBac* plasmid containing TetR-inducible Cas9 (Addgene #134247). After transfection, cells were selected with 800 μg/mL Geneticin for 7 days and single cell clones were generated using limiting dilution. To select the most optimal clone, Cas9 activity was evaluated by transducing each clone with the reporter lentiviral vector pXPR-011 (Addgene #59702, [31]), which expresses GFP and a sgRNA targeting *GFP*, followed by 4 days of 1 μg/mL doxycycline induction. GFP fluorescence was measured with high-throughput imaging at the single cell level to estimate the efficiency of the GFP gene knock-out.

After U2OS-iCas9 cell line was generated, the knockout of target genes was achieved by inducing Cas9 expression with 0.05 μg/mL doxycycline for at least 3 days.

### 4.3. sgRNA design and cloning

Either the CRISPick [32] or sgRNA Scorer [33] web-based software was used to design sgRNA CRISPR knock-out (KO) sequences targeting genes of interest as well as non-targeting controls. sgRNA sequences were individually synthesized (IDT) and independently cloned into lentiviral sgRNA expression vectors, either CROPseq-Guide-Puro (Addgene #86708) or CROPseq-puro-v2 (Addgene #127458), using NEBuilder HiFi DNA Assembly (NEB #E5520S) and BsmBI (NEB #0739S) restriction sites.

### 4.4. Lentivirus production, titration and transduction

Prior to lentiviral vector production, the corresponding targeting and non-targeting lentiviral sgRNA expression plasmids were mixed in equal quantities. HEK293T cells were seeded into 15 cm plates at a density of 8 million cells per plate. The following day, cells were transfected with pMD2.G (Addgene #12259), psPAX2 (Addgene #12260) and lentiviral expression plasmid (3:5:8 ratio by mass) using TransIT-Lenti transfection reagent (Mirus Bio #MIR6604) following manufacturer’s instructions. Lentiviral vector virions in the supernatant were harvested at 48 and 72 hours after transfection, gently mixed, and then filtered through a 0.45 μm filter. The lentiviral virions supernatant was further concentrated by adding 1:3 parts of Lenti-X concentrator (Takara #631232), incubated at 4°C for 4 hours and centrifuged at 1,500xg for 45 minutes at 4°C. The lentiviral virions pellet was resuspended in ice-cold PBS and stored at −80°C.

The lentiviral vector titer was individually determined for every cell line and sgRNA expression construct or mix of constructs. Cells were seeded at a density of 800,000 cells/well in a 12-well plate format using media supplemented with 8 μg/mL of polybrene (Biotechne #7711/10). Transduction was performed using viral vectors volumes ranging from 0 μL to 10 μL followed by centrifugation at 930xg for 2 hours at 33°C. After centrifugation, cells were incubated at 37°C for 24 hours prior to dividing them into media containing either 0 μg/mL or 2 μg/mL of puromycin (ThermoFisher Scientific #A1113803) for stable expression of the sgRNA. At 72 hours post antibiotic selection, cells from both conditions were counted and multiplicity of infection (MOI) was estimated and used to calculate average infectious units per microliter using the following formula: (MOI x seeding density)/viral volume.

Cells were transduced with viral sgRNA expressing vector pools in a 12-well plate format, using media supplemented with 8 μg/mL polybrene, an MOI of 0.1 and a minimum cell representation per sgRNA of 500x. Using a low MOI ensures that, on average, cells are transduced at most with one lentiviral copy per cell. Cells were then centrifuged at 930x g for 2 hours at 33°C followed by incubation at 37°C for 24 hours. Transduced cells were passaged into T-75 flasks containing media supplemented with 2 μg/ml puromycin. After 72 hours of selection, cells were expanded and used for further experiments.

### 4.5. *In situ* sequencing library preparation

Cells were seeded into either 384-well COC-bottom imaging plates (Revvity #6057308) or 6-well glass-bottom imaging plates (Cellvis #P06-1.5H-N) using doxycycline-free media. 48 hours after seeding, cells were fixed with 4% v/v paraformaldehyde (Electron Microscopy Sciences #15714) in PBS for 30 minutes, washed 3 times with PBS, and permeabilized with either 0.5% Triton X-100 in PBS or 70% v/v ethanol, depending on whether sequencing was combined with phenotype readout or not. When 70% ethanol was used for permeabilization, 75% volume exchanges with PBS containing 0.05% v/v Tweenx20 (PBS-T) were performed 6 times to prevent drying prior to 3 complete PBS-T washes. Reverse transcription mix (1x RevertAid RT buffer, 250 μM dNTPs (NEB #N0447L), 0.2 mg/mL recombinant albumin (NEB #B9200S), 1 μM LNA-modified RT primer, 0.4 U/μL RiboLock RNase inhibitor (ThermoFisher Scientific #EO0382), and 4.8 U/μL RevertAid H minus reverse transcriptase (ThermoFisher Scientific #EP0452)) was added to the samples followed by incubation at 37°C overnight. After reverse transcription, cells were washed 6 times with PBS-T, post-fixed with 3% paraformaldehyde and 0.1% glutaraldehyde in PBS for 30 minutes, and then washed 3 times with PBS-T. Samples were incubated in padlock probe and extension-ligation mix (1x Ampligase buffer, 0.4 U/μL RNase H (Enzymatics #Y9220L), 0.2 mg/mL recombinant albumin, 100 nM padlock probe, 0.02 U/μL TaqIT polymerase (Enzymatics #P7620L), 0.5 U/μL Ampligase (Lucigen #A3210K) and 50nM dNTPs) for 5 minutes at 37°C and 90 minutes at 45°C. After 3 PBS-T washes, rolling circle amplification (RCA) was performed by adding the corresponding reaction mix (1x Phi29 buffer, 250 μM dNTPs, 0.2 mg/mL recombinant albumin, 5% glycerol and 1 U/μL Phi29 DNA polymerase (ThermoFisher Scientific #EP0091) and incubating samples at 30^°^C overnight.

When sequencing-by-synthesis (SBS) reactions were not performed, rolling circle amplicons (rollonies) were detected by incubating samples with 100 nM Alexa-647 labelled fluorescent sequencing primer (IDT), 2X SSC, and 10% formamide for 30 minutes at 37°C. After 3 PBS-T washes, cells were incubated in imaging solution containing 200 ng/mL of DAPI (4′,6-diamidino-2-phenylindole) in 2X SSC.

### 4.6. Immunofluorescence

For immunostaining experiments, cells were seeded into either 384-well COC-bottom plates or 6-well glass-bottom plates at least 24 hours before fixation. Cells were fixed with 4% v/v paraformaldehyde in PBS for 30 minutes and permeabilized using 0.5% Triton X-100 in PBS for another 30 minutes. After permeabilization, cells were washed 3 times with PBS-T and incubated in blocking buffer containing 5% Bovine Serum Albumin (BSA) in PBS-T for 30 minutes. Mouse-anti-LMNA/C antibody (Santa Cruz Biotechnology #B0719, 1:1,000 dilution in blocking solution) was used to stain the cells for 1 hour at room temperature followed by 3 10-minutes washes with PBS-T and incubation with anti-mouse Alexa-488 secondary antibody (Cell Signaling Technology #4408S, 1:2,000 dilution in blocking buffer) for another hour at room temperature. After 3 10-minutes washes with PBS-T, a solution containing 200 ng/mL DAPI in 2X SSC was used for imaging.

When immunofluorescence was combined with ISS, the protocol was started at the blocking step and after ISS post-fixation.

### 4.7. Telomeric native FISH (nFISH)

When telomeric nFISH was not paired with ISS, cells were seeded into either a 384- or a 6-well plate at least 24 hours prior to fixation. Cells were fixed using 4% v/v paraformaldehyde in PBS for 30 minutes, washed 3 times with PBS, and permeabilized with 0.5% Triton X-100 in PBS. After 3 PBS-T washes, cells were incubated in blocking buffer containing 1 mg/ml BSA, 3% goat serum, 0.1% Triton X-100, 1 mM EDTA pH 8, and 500 μg/mL RNase A (ThermoFisher Scientific #12091021) for 1 hour at 37°C. Cells were then washed 3 times with PBS-T and consecutively dehydrated with increasing concentrations of EtOH (75%, 95%, and 100%) for 5 minutes at a time. Samples were air dried before adding hybridization solution. Hybridization solution, incubation time and washes were different depending on telomeric probe used. Cells were incubated in PNA-probe mix (70% deionized formamide (Invitrogen #AM9344), 0.5% blocking reagent (Roche #11096176001), 10mM Tris-HCl and 1:1000 PNA probe (PNABio #F1006)) for 1 hour at room temperature followed by two 15-minute incubations in washing buffer 1 (70% deionized formamide, 10mM Tris-HCl) and 3 PBS washes. Alternatively, cells were incubated in unmodified probe mix (50% deionized formamide, 0.2% blocking reagent, 2X SSC, 4:1000 unmodified probe) for 40 hours at room temperature, followed by two 15-minutes incubations in washing buffer 2 (0.1% Tween20, 2X SSC) and 3 0.2X SSC washes. A solution containing 200ng/mL DAPI in 2X SSC was used for imaging.

When telomeric nFISH was paired with ISS, the protocol was started at the blocking step either after ISS post-fixation or RCA.

### 4.8. Small molecule inhibitors

ALT activity was further enhanced in U2OS cells by treating them with 10 μM of E804-20 [16], a chemical inhibitor of the Tousled Like Kinase 1 and 2 proteins (TLK1/2), for a total of 6h before cell fixation. E804-20 was purchased from ChemMaster International.

### 4.9. Sequencing-by-synthesis (SBS)

The sgRNA sequences copied in the amplicons were read out using SBS reagents from the Illumina MiSeq 500 cycle Nano kit (Illumina #MS-103-1003). Cells were incubated in a mix containing 1 μM sequencing primer, 2X SSC and 10% formamide for 30 minutes at 37°C. Cells were then washed 3 times with incorporation buffer (Nano kit PR2) and incubated with incorporation mix (Nano kit reagent 1) for 5 minutes at 60°C. The incorporation mix was gradually washout over six serial dilutions with PR2 buffer and samples were further washed five times with fresh PR2 at 60°C for 5 minutes each. After these washes, a solution containing 200 ng/mL DAPI in 2X SSC was used for imaging. Following each imaging cycle, dye terminators were removed by incubating cells with cleavage mix (Nano kit reagent 4) for 8 minutes at 60°C and by then washing cells with PR2 six times for 1 minute at 60°C each.

### 4.10. Fluorescence microscopy

A Nikon Ti-2 Eclipse inverted epifluorescence microscope with automated stage control, a 16-bit sCMOS camera, and hardware autofocus was used to acquire phenotypic and ISS images in 6-well plates. All hardware was controlled using NIS-Elements AR software, and a LED light engine (Spectra III, Lumencor, 390/440/475/510/555/575/635/747 nm excitation wavelengths) was used for fluorescence illumination in addition to a penta excitation dichroic mirror (353-403 nm, 462-486 nm, 554-556 nm, 627-643 nm, 722-748 nm). The lamin A phenotype was imaged using a 20X air 0.95 NA CFI Apo Lamba S objective and the following Semrock filters and exposure times: excitation 488 nm with 30% power, FF01-515/30-32 emission and 30 ms. The nFISH phenotype was imaged using a 40X water (NA 1.25) CFI Apo Lambda objective and the same optical configuration but with 300 ms exposure time. ISS images were acquired using a 10X air (NA 0.45) CFI Plan Apo Lambda objective and the following Semrock filters and exposure times for each channel: DAPI (excitation 365nm with 24% power, FF01-432/36-32 emission and 100ms), MiSeq G (excitation 561nm with 65% power, FF01-515/30-32 emission and 300ms), MiSeq T (excitation 561nm with 10% power, FF02-615/20-25 emission and 200ms), MiSeq A (excitation 640nm with 5% power, FF01-676/29-25 emission and 100ms) and Miseq C (excitation 640nm with 90% power, FF02-809/81-32 emission and 200ms).

A dual spinning disk high-throughput confocal microscope (Yokogawa CV8000) with a 405/488/561/640 nm excitation dichroic mirror and two 16-bit sCMOS cameras with binning set to 2 was used to acquire phenotypic and rollonies images in 384-well plates. Emission bandpass filters were used for each channel: 445/45, 525/50, 600/37, and 676/29 nm. Rollonies and lamin A images were acquired using a 20X air (NA 0.75) objective, and nFISH images were acquired using a 40X water (NA 1.1) objective.

### 4.11. Image analysis (phenotyping + ISS)

To extract sgRNA guides from ISS data, we adapted and further developed a comprehensive image analysis pipeline based on a previously described approach [11]. We first segmented nuclei from images from the DAPI channel in each SBS cycle using the Cellpose deep learning model [34,35] to define single cells, followed by cytoplasm segmentation by calculating the median across all 4 non-DAPI channels as input for watershed algorithm with the nuclei regions of interest (ROIs) as seeds. Next, we aligned images across all 12 sequencing cycles using cross-correlation of the DAPI channel to ensure spatial consistency and accuracy, mitigating potential translational shifts or distortions during imaging. The image alignment step ensures that all sequencing channels remain properly registered relative to the nuclei, which serve as a stable reference across cycles. To detect ISS spot-like signals, we computed an image where each pixel represents pixel-level intensity standard deviation (SD) across all 12 SBS cycles for each SBS channel, and then averaged the SD images across all 4 SBS channels. High values in the averaged SD image represents areas with large cycle-to-cycle fluorescence changes, which are expected to be areas where ISS spot-like signals are located. Finally, we performed base calling by quantifying fluorescence intensities at the detected spots for each cycle, generating 12-nt sgRNA reads for each detected signal. To improve accuracy, a median correction was applied to the raw base signal using a transformation matrix derived from the median fluorescence intensities across the four nucleotide channels (G, C, A, T). The corrected signals were then used for base calling, where nucleotide identities were assigned to each spot in a cycle based on the highest corrected fluorescence intensity. This process was repeated across all 12 sequencing cycles, resulting in a complete 12-nt sgRNA sequence for each detected spot.

For each cell, we took into account all the ISS reads overlapping with its cell body, counted the frequency reads for each sgRNA sequence, selecting the sequence with the highest read count as the representative sgRNA. To identify the dominant barcodes in each cell, the top two most frequently occurring barcodes were selected, along with their corresponding counts, to ensure accurate representation of the dominant barcodes. Finally, we mapped the identified sgRNA sequences to their target genes using a predefined codebook using a Levenshtein distance of 0 to ensure exact matches.

We employed a global coordinate-based approach to match the ISS data acquired from images at 10X magnification with phenotype data acquired from images at either 20X magnification (LMNA IF) or 40X magnification (nFISH). This utilized the coordinates embedded within each field of view (FOV) in ND2 files to establish a common coordinate system. We then calculated the offset between the 10X ISS and 20X/40X phenotype images by determining the displacement between overlapping FOVs, where the 20X/40X FOV was nested within the corresponding 10X FOV, enabling accurate mapping of sgRNA information corresponding to cells in the higher-resolution phenotype images. Morphological properties, including perimeter, area, eccentricity, and solidity, were used to verify accurate correspondence between cells. A matching threshold of 10% was applied to ensure that only cells with morphological parameters within this threshold were considered valid matches. This allowed us to determine the perturbed gene for each cell, facilitating downstream analysis of gene perturbation effects.

The entire pipeline was designed for high-throughput processing of the images. We optimized the code for parallel processing, leveraging the cluster’s multi-node capabilities and NVIDIA A100 GPUs for efficient handling of large datasets. We used several scientific Python modules for the implementation of this analysis and data processing. NumPy facilitated numerical computations for handling arrays and matrices, while Matplotlib generated plots and visualizations. Pandas managed tabular data, and Cellpose, along with scikit-image, enabled segmentation and advanced image processing. Pillow handled basic image manipulations, and Threading enhanced computational efficiency by parallelizing tasks. Editdistance computed Levenshtein distances for barcode sequence matching, and SciPy performed statistical and mathematical operations. The code for the analysis can be found at https://github.com/CBIIT/HITIF-NFISH-ISS.

### 4.12. Statistics

Statistical significance for integrated nFISH intensity between cells perturbed with every sgRNA and control cells was calculated using the non-parametric Mann-Whitney U test and the Bonferroni correction was applied to adjust significance levels for each test. Other two group comparison statistics were also performed using the non-parametric Mann-Whitney U test. For every experimental condition a minimum of 3,000 cells and three technical replicates were analyzed.

## 5. Acknowledgements

We would like to thank Tom Misteli for critical reading of the manuscript. HiTIF and the Lazzerini Denchi lab were supported by the Intramural Program of the National Cancer Institute (Projects 1-ZIC-BC011567-11 and ZIABC011811816). This work utilized the computational resources of the NIH HPC Biowulf cluster (https://hpc.nih.gov).

